# Loss of pH switch unique to SARS-CoV2 supports unfamiliar virus pathology

**DOI:** 10.1101/2020.06.16.155457

**Authors:** Kristina A. Paris, Ulises Santiago, Carlos J. Camacho

**Affiliations:** Department of Computational and Systems Biology, University of Pittsburgh, Pittsburgh, PA, 15260, USA

**Keywords:** SARS-CoV, SARS-CoV2, binding free energy, pH, shear flow, infectivity, incubation, shedding, transmission rate, viral entry

## Abstract

Cell surface receptor engagement is a critical aspect of viral infection. This paper compares the dynamics of virus-receptor interactions for SARS-CoV (CoV1) and CoV2. At low (endosomal) pH, the binding free energy landscape of CoV1 and CoV2 interactions with the angiotensin-converting enzyme 2 (ACE2) receptor is almost the same. However, at neutral pH the landscape is different due to the loss of a pH-switch (His445Lys) in the receptor binding domain (RBD) of CoV2 relative to CoV1. Namely, CoV1 stabilizes a transition state above the bound state. In situations where small external strains are applied by, say, shear flow in the respiratory system, the off rate of the viral particle is enhanced. As a result, CoV1 virions are expected to detach from cell surfaces in time scales that are much faster than the time needed for other receptors to reach out and stabilize virus attachment. On the other hand, the loss of this pH-switch, which sequence alignments show is unique to CoV2, eliminates the transition state and allows the virus to stay bound to the ACE2 receptor for time scales compatible with the recruitment of additional ACE2 receptors diffusing in the cell membrane. This has important implications for viral infection and its pathology. CoV1 does not trigger high infectivity in the nasal area because it either rapidly drifts down the respiratory tract or is exhaled. By contrast, this novel mutation in CoV2 should not only retain the infection in the nasal cavity until ACE2-rich cells are sufficiently depleted, but also require fewer particles for infection. This mechanism explains observed longer incubation times, extended period of viral shedding, and higher rate of transmission. These considerations governing viral entry suggest that number of ACE2-rich cells in human nasal mucosa, which should be significantly smaller for children (and females relative to males), should also correlate with onset of viral load that could be a determinant of higher virus susceptibility. Critical implications for the development of new vaccines to combat current and future pandemics that, like SARS-CoV2, export evolutionarily successful strains via higher transmission rates by viral retention in nasal epithelium are also discussed.

## Introduction

Although accurate assessments are still evolving, reports from the world health organization indicate that infection with SARS-CoV-2 (CoV2) is significantly different relative to infection with previous respiratory viruses. The biggest distinctions are the longer incubation period and increased viral shedding (both likely responsible for higher rates of transmission), strong correlation of infected-fatality rates (IFR) with age and comorbidities, higher IFR for males relative to females, and minimal impact in children. As much as 40% of deaths from CoV2 relate to cardiovascular complications (Akhmerov and Marban, 2020), while cellular and animal models have also revealed inappropriate inflammatory responses along with high chemokine production (Blanco-Melo et al., 2020). Using thermodynamic, kinetic and molecular modeling, we explored potential reasons for these CoV2-unique aspects and aimed to identify determinants of its complex pathology. While it stands to reason that some of the answers may be found in the complex genomic changes triggered by the virus in cells, tissues, and organs, longer incubation and infectivity time scales also suggest that differences could have a biophysical origin.

Work on SARS-CoV (CoV1) has already determined that the virus enters cells via receptor-mediated endocytosis in a pH-dependent manner (Wang et al., 2008) that is characterized by co-translocation of the viral spike glycoprotein and its specific functional receptor, the angiotensin-converting enzyme 2 (ACE2), from the cell surface to early endosomes. Key steps that control the fate of the virus in the early and late endosome are driven in part by lowering the pH from 6.5-to-6.0 and from 6.0-to-5.0 (Bui et al., 1996), respectively; exposure to low pH triggers a spike catalyzed fusion between the viral and endosomal membranes followed by viral genome release. The cell machinery is then hijacked to replicate and assemble new virus particles that eventually exit the cell, primarily through budding. Broadly speaking, this is the same pH-dependent endocytic path followed by the influenza virus (Qin et al., 2019; Yamauchi, 2019), and likely all other coronaviruses (including MERS-CoV).

While infections by CoV1 and CoV2 are mediated by the ACE2 receptor, MERS-CoV (MERS) gains entry to cells through DPP4 (Raj et al., 2013). Figure 1 shows structures for the receptor-binding domains (RBD) in complex with their receptors for all three viruses. Surprisingly and likely significantly, while the RBD of CoV1 (Li et al., 2005) and MERS (Wang et al., 2013) have one Histidine on opposite ends of their binding interface, CoV2 (Wang et al., 2020) does not have His residues in this domain. As we have been able to determine to date, all known strains of CoV2 have mutated away their last His residue that is still present in the RBD of CoV1/MERS and related zoonotic viruses (see below). Because cell surface receptor engagement is a critical aspect of viral infection and life cycle, and sensing pH is relevant for both viral replication and regulation of Histidine protonation, we set to decipher the mechanistic role of the remaining His residue that distinguishes the RBDs of CoV1 and CoV2.

**Figure 1.**
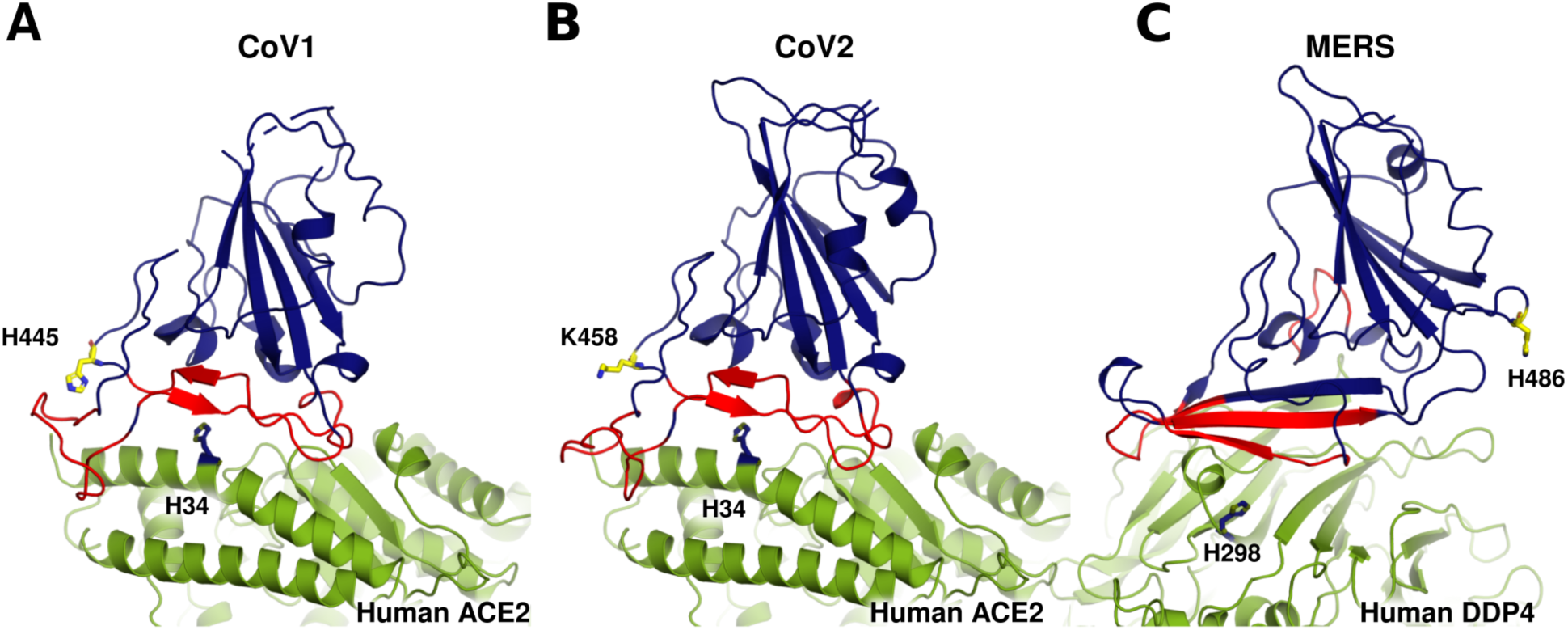
Deletion of Histidine residue in receptor binding domain (RBD) of CoV2. Co-crystal structures of the receptor binding domain (RBD) and the receptor for (A) SARS-Cov (PDB 2AJF; CoV1), (B) SARS-CoV2 (PDB 6LZG; CoV2) and (C) MERS-CoV (PDB 4L72; MERS). RBD of the viruses is in blue with binding interface in red. Receptor of Cov1 and Cov2 is ACE2, and of MERS is DPP4. RBD’s of Cov1 and MERS have one histidine (H), H445 and H486, respectively, shown in yellow. ACE2 and DPP4 have several histidine residues, only the closest to the binding interface is shown in blue.

## Results

### pH-switch

At low (endosomal) pH, the binding free energy landscape of CoV1 and CoV2 interactions with their ACE2 receptor is almost the same. This is important because low pH is critical for the activation of the spike fusion in late endosomes (Martin and Helenius, 1991). In particular, His34 in ACE2 (Fig. 1), located at the core of the RBD binding interface, should play a key role in this process. Indeed, we predict based on the co-crystal structures that His34^+^ should readily form a hydrogen bond network that stabilizes the RBD/ACE2 complex in both CoV1 and CoV2 (Fig. 2). Thus, loss of the pH-switch in CoV2 has no impact on the low pH bound conformation with ACE2.

**Figure 2.**
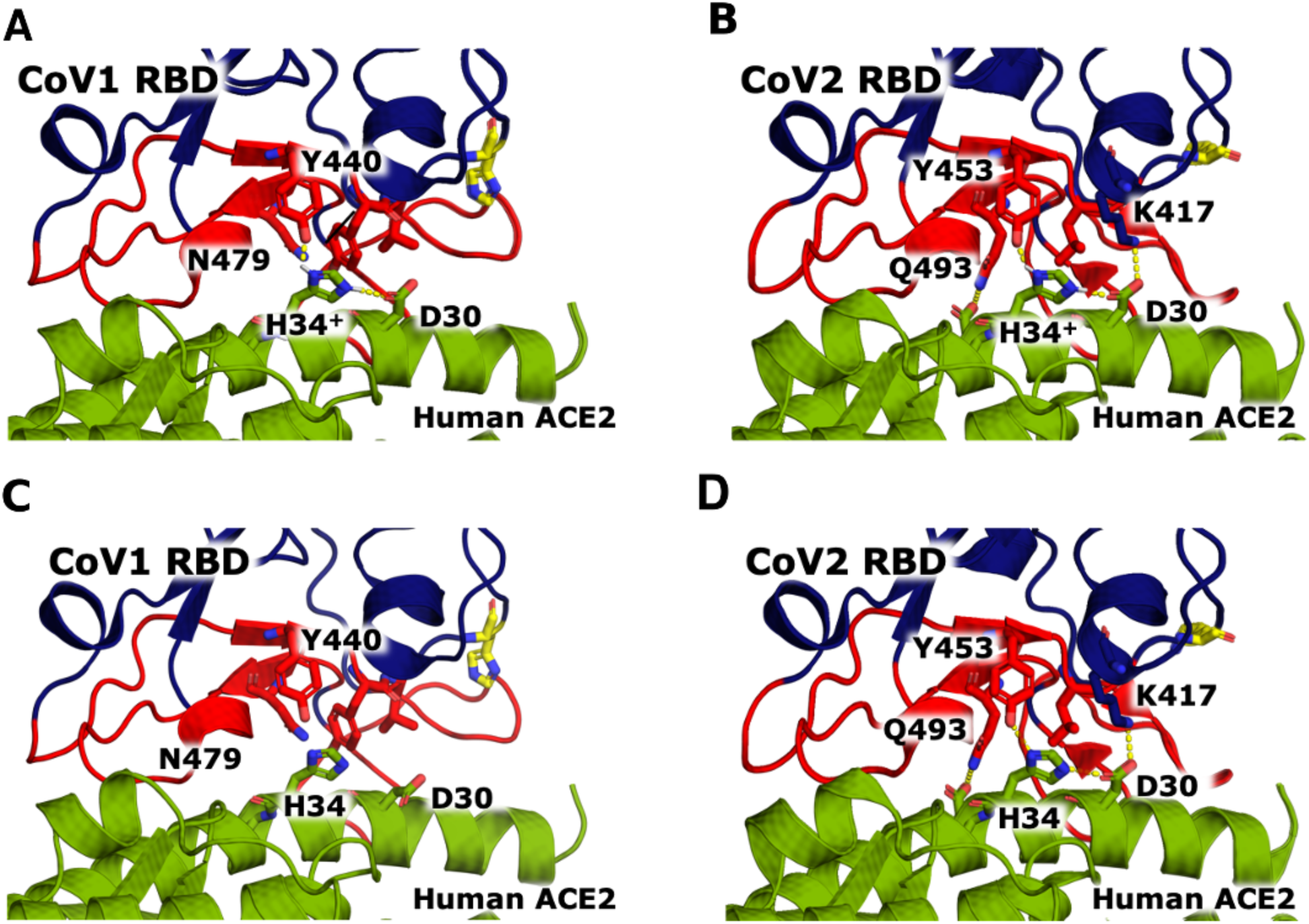
Protonation of the Human ACE2 H34 further stabilizes binding interface. Protonated state of H34^+^ is predicted to form stable H-bond network with D30 in ACE2, and Y440 and Y453 in (A) CoV1 and (B) CoV2, respectively. H-bond network is based on rotamers already observed in co-crystals of the unprotonated forms: (C) PDB 2AJF for CoV1, and, (D) PDB 6LZG for CoV2 (see also PDB 6M0J). Unprotonated co-crystal structures of CoV2 assigned H34 with a NδH making a bond with backbone oxygen of D30, which is already making a bond in the α-helix. Our MDs indicate that even in the unprotonated form the rotamer should be rotated 180°having Nδ and Nε more readily interacting with D30 and Y453 (as shown in panel D).

On the other hand, at neutral pH the landscape is different due to the loss of the pH-switch (His445Lys) in the RBD of the spike protein of CoV2 relative to CoV1. We studied this loss by performing three independent unconstrained molecular dynamics simulations (MDs) of CoV1 PDB 2DD8 (Prabakaran et al., 2006) and CoV2 from the receptor (apo) in PDB 6LZG at both physiological and low pH ∼ 5.0 conditions (See Methods in Supplementary Information). In practice, lowering the pH protonates the His residue from neutral to positively charged (His^+^) by the addition of an extra hydrogen. Of note, although not studied here, the pH-switch in the RBD of MERS has previously been observed in pH-dependent crystal structures (Zhang et al., 2018).

MDs revealed two distinct regions in the binding interface of CoV1: (a) a loop motif (F_460_SPDGKPCTPPALNCY_475_) that for His and His^+^ shows mostly not-bound- and bound-like conformations, respectively; and, (b) the rest of the binding interface that consistently adopts bound-like conformations that are independent of pH (Fig. 3A-C). Figure 3A-B shows the dominant clusters observed in the His and His^+^ simulation of CoV1, as well as the pH-independent simulation of CoV2. Figure 3C-D shows detailed analyses of the corresponding root-mean-square-deviations (RMSDs) of these two regions relative to the co-crystals as a function of time (Additional MDs are shown in Fig. S1). The plots include the equilibration time (between 100-200 ns) in order to emphasize that the distinct trajectories were not biased by different initial conformations. Remarkably, deletion of the pH switch in CoV2 that mutates the last Histidine, His445, in CoV1 by Lys458 in CoV2 generates a motif, which includes T_470_EIYQAG_476_, that yields almost exclusively bound-like conformations (Fig. 3D)—i.e., CoV2 is always ready to bind ACE2. It is interesting to note that the effect of both Lys458 in CoV2 and His^+^ in Cov1 (Fig. 3C-D) is to stabilize the bound-like state.

**Figure 3.**
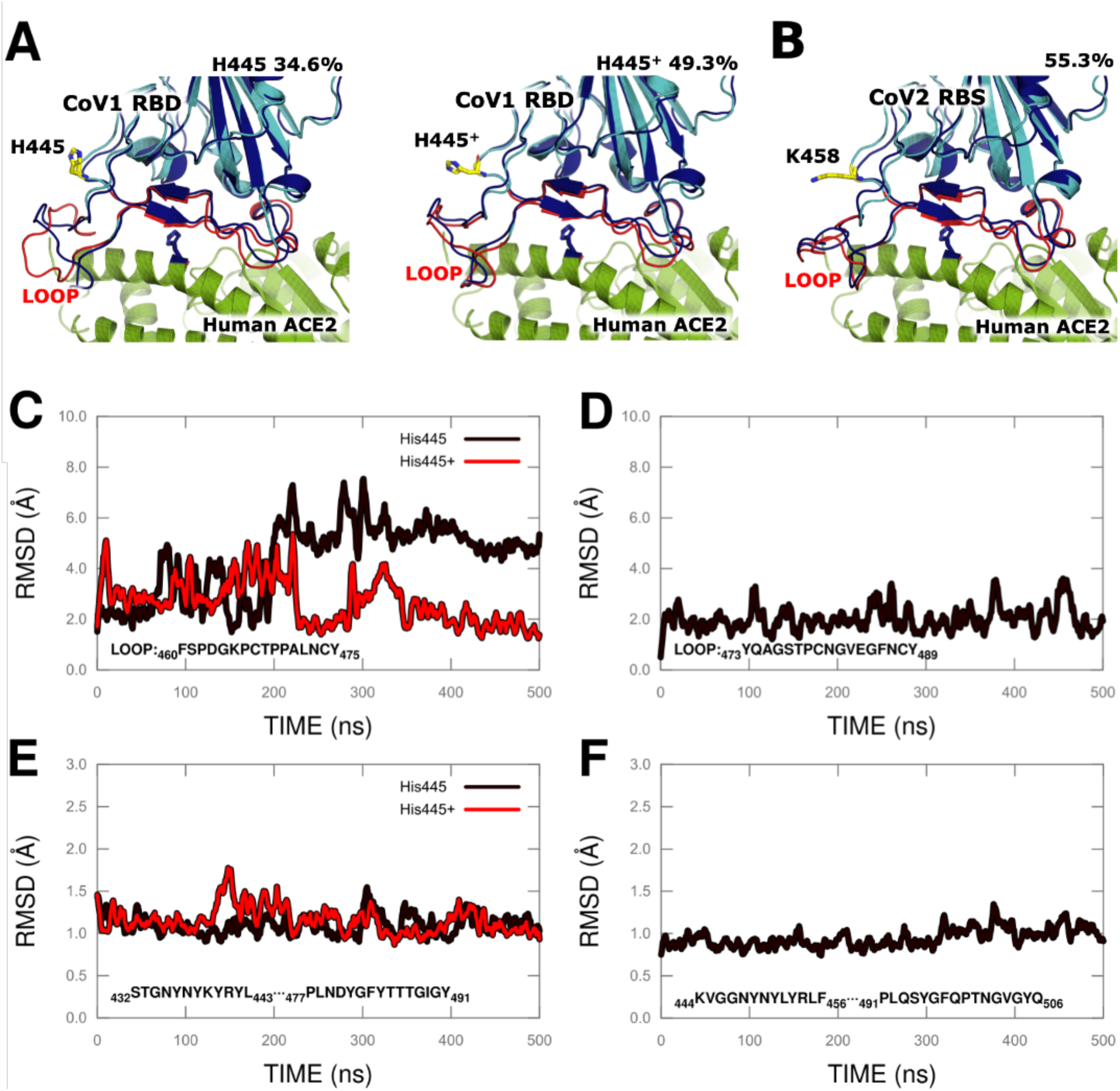
Role of pH switch in CoV1 relative to CoV2. (A) and (B) show the same co-crystals as in Fig. 1 superimposed with centroid of largest 1.8 Å RMSD cluster of conformations of pH-switching loop from MDs shown in (C) for CoV1 and (D) for CoV2. Also indicated is the size of the corresponding cluster relative to simulation time. (C) Root-mean-squared-deviation (RMSD) of 16 amino acid loop as a function of time that switches between not-bound-like (∼ 5.8 Å) to bound-like (∼ 1.9 Å) relative to co-crystal (PDB 2AJF), for His and His^+^, respectively; (D) Same analysis for CoV2 homologous loop shows most conformations under 2 Å RMSD relative to PDB 6LZG. Binding interface, not including loop, stays in a bound-like conformation for 100% of the simulation time for both (E) CoV1 and (F) CoV2.

Implications of these findings in the binding free energy landscape are sketched in Fig. 4. Namely, at low (late endosomal) pH, the landscape of CoV1 and CoV2 interactions with the ACE2 receptor are very similar. However, at neutral pH the landscape is different due to the loss of the pH switch in the RBD of CoV2 relative to CoV1. Specifically, the not-bound-like conformations of the pH-dependent loop in CoV1 stabilizes a higher free energy transition state, whereas the persistent bound-like behavior of CoV2 yields a much tighter bond.

**Figure 4.**
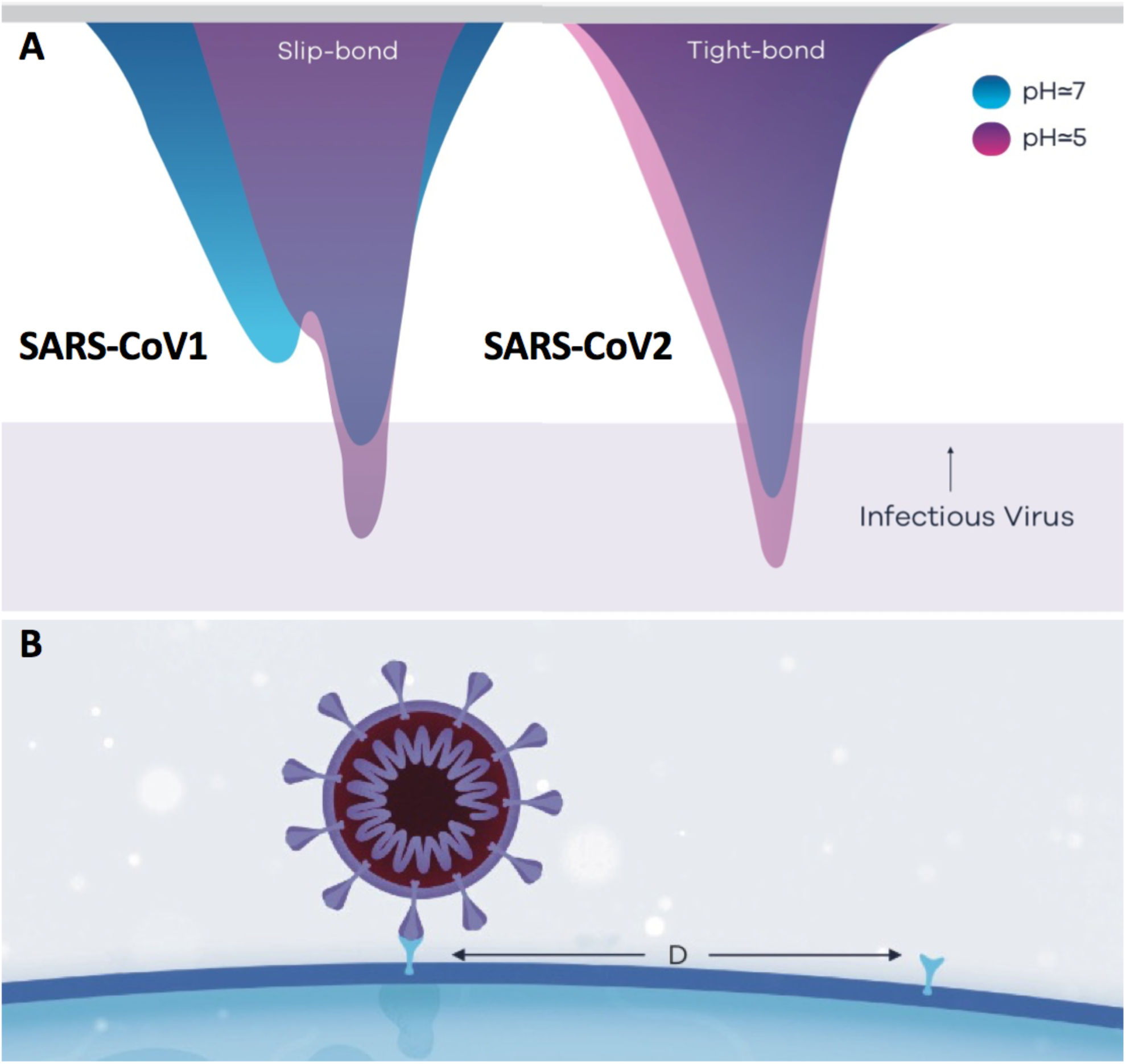
Free energy binding landscapes of RBD of SARS-CoV1 and SARS-CoV2 and ACE2 receptor. (A) At physiological pH, CoV1 has a pH-dependent transition state that significantly weakens the RBD/ACE2 bond when small external strains are applied. Although CoV2 conformations do not depend on pH, protonation of ACE2 further stabilizes the complexes as depicted for pH ≈ 5. (B) Cartoon of virus attached to one cell receptor in respiratory airways, recruitment of a second receptor depends on balance between dwelling time of the bond and diffusion time scale by a distance D of protein in cell membrane, where D is average distance between receptors.

### Dynamics of virus-receptor interactions for CoV1 and CoV2

What are the implications of this transition state found in CoV1 but not in CoV2 at neutral pH? To answer this question, we need to consider that viral particles in the respiratory system are under small external strains from, e.g., shear flow in the respiratory airways, when engaging cell surface receptors.

The cell mechanics of these interactions can be described by applying the reaction-limited kinetics of membrane-to-surface adhesion and detachment first envisioned by Dembo et al (Dembo et al., 1988). Specifically, one can write the free energy of the bound-state (bs) Δ*G*_*bs*_ under a tensile force as a function of the cell-cell gap width L, a constant binding free energy 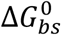 and a “spring” energy such that

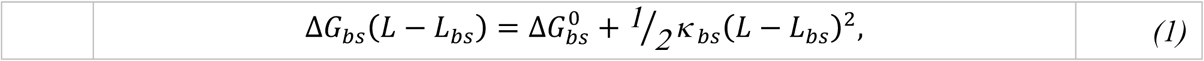

where *L*_*bs*_ is the equilibrium length for the bonded state. A similar equation holds for the free energy of the transition state (ts) Δ*G*_*ts*_ at the same cell-cell gap,

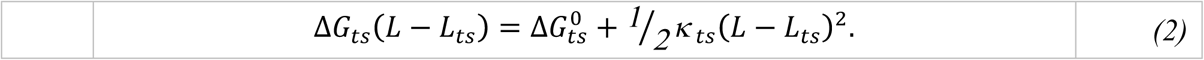

This treatment assumes equilibrium between bonded and de-bonded states, so there must be a very slow “ramp” rate for the force of pulling or pushing. If these conditions apply, the equilibrium constant for bond formation can be written as

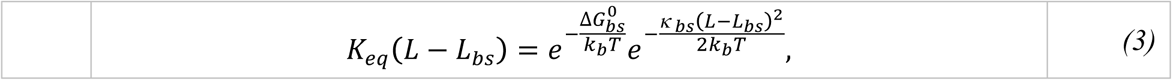

where *k*_*b*_*T* is the Boltzmann factor. Note that

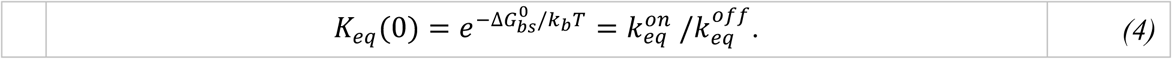

According to Arrhenius theory, the de-bonding rate constant at a given gap-width can then be written as

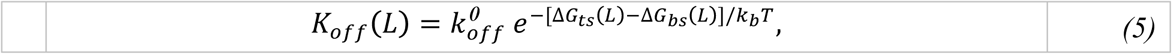

The mechanical or structural basis of the RBD/receptor interactions in Fig. S2 can be characterized as a “door-knob” type junction across the gap, as opposed to a gripping or fish-hook bound state. The knob interactions with the receptor entailed two characteristic patches, a large bound-like domain and the smaller switching loop (Fig. 3), which one could model with spring constants κ _*bl*_ and κ _*loop*_, respectively (see red and yellow surfaces in Fig. S2). Then, the elastic constant of the bound- and transition-state can be written as κ _*bs*_ ≈ κ _*bl*_ + κ _*loop*_ and κ _*ts*_ ≈ κ _*bl*_, respectively, with the equilibrium rest lengths being essentially the same, i.e., L_*bs*_ = *L*_*ts*_ = L_*0*_. Thus, virus detachment to the transition state corresponds to an ideal case of the theory, where the only allowed change between the bonded and transition state is in the spring constants, such that

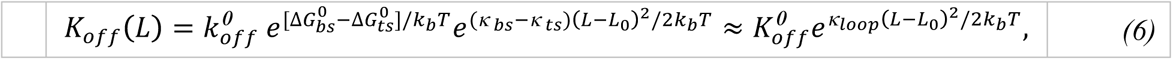

Where

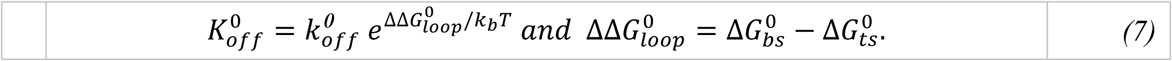

It is clear that the spring approximation should only apply for small deformations. In general, the springs across the gap undergo a “twisting” motion around its long-axis to reach the transition state from the bonded state. The motion will increase “tightness” of the spring if (κ _*bs*_ − κ _*ts*_) < 0, which defines a catch-bond. Here, however, CoV1 always loosens tightness, (κ _*bs*_ − κ _*ts*_) = κ _*loop*_ > 0, corresponding to a typical slip-bond whose lifetime is shortened by tensile forces acting in the bond.

### Free energy landscapes and estimates of bond detachment for CoV1/CoV2 and ACE2

We use the *FastContact* server (Champ and Camacho, 2007) to compute the electrostatic 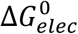 and desolvation 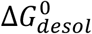 binding free energies of the bound and transition states using co-crystal structures and chimeras that incorporate changes triggered by low pH structures. Entropies coupled to the unbound state could be somewhat higher for CoV1 relative to CoV2 due to the larger conformational entropy associated with the switching loop in Fig. 3. Other error bars are correlated since interactions are very similar such that ΔΔ*G*′*s* have an error bar of ± 1 *kcal*/*mol*. Absolute free energies need to account for size-dependent configurational and vibrational entropy changes upon binding, which for high affinity protein-protein complexes have been estimated to be anywhere between 5-to-15 kcal/mol. However, for flat and rather superficial complexes such as those here (Fig. S2), the entropy loss could be much lower. Finally, the pre-exponential factor 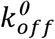 (Eq. 5) in the absence of a transition state and at equilibrium is exactly 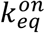 at 1 M concentration, which for diffusion-limited association can be approximated by 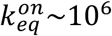s^-1^ (Camacho et al., 2000). Table 1 lists 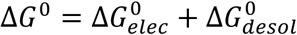 and the baseline rate of detachment for the RBD/ACE2 bond under tension 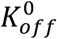 to the (a) transition state (ts) of CoV1, and full detachment of the (b) transition (ts) of CoV1 and (c) CoV2

**Table 1.**
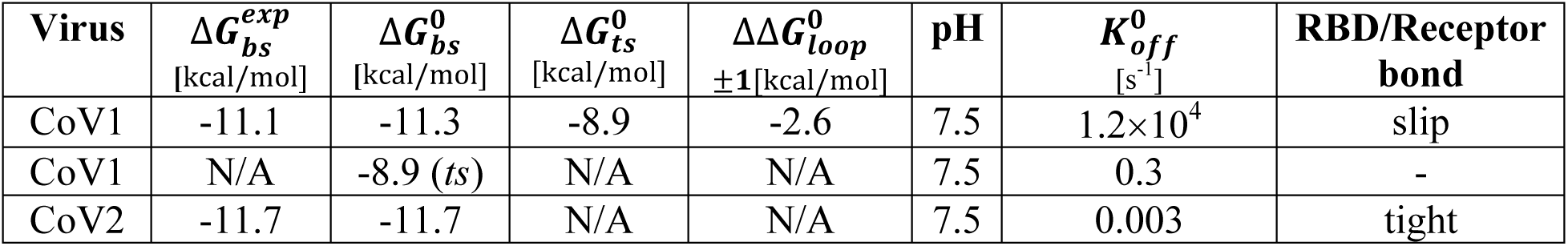
Free energy estimates and baseline rate of detachment of RBD/ACE2 complex. Estimates of bound-state (bs) are computed by *FastContact* server and co-crystals shown in Figs 1 and 2 (neutral pH). Free energy of transition state in CoV1 was computed by eliminating the contribution of the switching loop 460-475 that acquires not-bound-like conformations from *FastContact* calculation (Fig. 3A-C). Experimental equilibrium binding free energy is from (Walls et al., 2020).

| Virus | $\Delta G_{bs}^{exp}$<br>[kcal/mol] | $\Delta G_{bs}^0$<br>[kcal/mol] | $\Delta G_{ts}^0$<br>[kcal/mol] | $\Delta \Delta G_{loop}^0$<br>$\pm 1$[kcal/mol] | pH | $K_{off}^0$<br>[s<sup>-1</sup>] | RBD/Receptor<br>bond |
| --- | --- | --- | --- | --- | --- | --- | --- |
| CoV1 | -11.1 | -11.3 | -8.9 | -2.6 | 7.5 | $1.2 \times 10^4$ | slip |
| CoV1 | N/A | -8.9 (ts) | N/A | N/A | 7.5 | 0.3 | - |
| CoV2 | -11.7 | -11.7 | N/A | N/A | 7.5 | 0.003 | tight |

Free energy estimates of bound complexes are fully consistent with experimental data (Walls et al., 2020); alternative measurements have suggested a 10 fold weaker binding (Shang et al., 2020). These differences will only re-scale 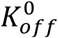 by a factor of 10 but will not be significant to our conclusions. The key observation is that CoV1 stabilizes a transition state by about 2.6 *kcal*/*mol* above the bound state. As a result, small external strains applied by, say, shear flow in the respiratory system enhance the off rate of the viral particle as shown in Eq. 6. Thus, CoV1 virions are expected to detach from cell surfaces in faster time scales. These binding free energy estimates are depicted in the landscapes in Fig. 4. Only at physiological pH should the landscape of CoV1 display a pH-dependent transition state. Other bonds are expected to break in an all-or-none type of transition.

### Optimal dwelling times and endocytosis

In principle, the pH-switch in CoV1 could provide a natural mechanism to optimize virus internalization. Namely, CoV1 is expected to “bounce around” cell surfaces many times before cell entry. If the density of receptors is high enough, a “stick-and-slip” approach could be an efficient mechanism to find clusters of receptors randomly distributed on the cell surface. On the other hand, if only a small number of cell surface receptors are available, then receptor diffusion will be the limiting step to accrue the critical number of receptors needed for endocytosis, and longer RBD/ACE2 dwelling times will be required. Of note, tighter binding to ACE2 would also make it easier for a smaller number of CoV2 particles to establish an initial foothold in the respiratory system compared to weak binding where particles could be exhaled out.

Broad estimates of “high” concentration, e.g., in the range of 1000-to-10,000 receptors, yield an average separation between receptors *D*∼0.6 − 0.2 *μm* (see Fig. 4B) that is larger than the diameter of the virus ∼0.12 *μm*. Thus, after attachment of the first spike to its receptor, recruitment of a second receptor to stabilized virus attachment will be limited by other receptors circulating in the cell membrane. Lateral protein diffusion in cell membranes is length-scale dependent, varying between 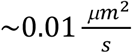 (Spendier et al., 2012) and 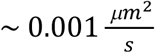 (Kusumi et al., 2005) for 40-100 nm and >100 nm, respectively. Thus, diffusion time scales to bring two receptors into close proximity for the above length scales are ∼200 − 5 *s*.

It is noteworthy that the number of surface receptors in cells have an upper limit of about 20,000, which in the respiratory airways could limit infectivity to dwelling times of about 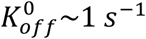 (or *K*_12_∼10^6^ *M*^−1^) based on *D*∼0.15 *μm* (Fig. 4).

## Discussion

### Viral infection and its pathology

The lifetime of CoV1 RBD/ACE2 bonds at physiological pH (∼3 s) is marginally short-lived for efficiently triggering endocytosis, even at high ACE2 receptor concentrations. As a result, CoV1 virions are expected to detach from cell surfaces in time scales that are much faster than the time needed for other receptors to reach out and stabilize virus attachment. And for human nasal goblet cells, it will be significantly worse since, after each bounce, particles will be biased by gravity to either diffuse down the respiratory tract or be exhaled, where they will not find significant amounts of ACE2 receptors until reaching lung alveolar epithelial cells (Hamming et al., 2004). On the other hand, deletion of the pH switch allows CoV2 to have RBD/ACE2 bonds with dwelling times of about ∼ 300 s, commensurate with the diffusion time scales needed to recruit enough ACE2 receptors to trigger endocytosis. This mechanism implies that, for the most part, CoV1 will not co-localize in the nasal cavity. This prediction is consistent with CoV1 being mainly a lower respiratory tract disease, causing complications that include acute respiratory distress (Ding et al., 2003; Hamming et al., 2004).

Viral replication in human mucous gland cells will release viruses back into the same area where they can infect new cells until the supply of ACE2 receptors is depleted below the critical threshold needed for binding and internalization. This process will trap viral particles in the upper respiratory tract, naturally leading to longer incubation times. Similarly, accumulation of viral particles in the nasal mucosa will lead to extended periods of viral shedding. Of note, since viral transit to the lower respiratory tract will be significantly slower for CoV2 relative to CoV1, this period of higher infectivity rates could be for the most part mediated by asymptomatic individuals.

Based on our findings, incubation times should correlate with the number of ACE2-rich cells in the nasal area. It is important to note that children do not have well developed sinuses until adolescence (Henson et al., 2020). Thus, large areas for viral replication will not be available in children, resulting in shorter incubation times due to the faster diffusion down to the lower respiratory tract. Something similar could apply to females who have smaller nasal cavities relative to males (Samolinski et al., 2007). Shorter times in the nasal cavity would lead to a lower viral load in the upper airways and could explain the lower transmission and milder symptoms that are observed in children, as well as the lower IFR in adult women relative to men.

Our proposed mechanism is also consistent with reported loss of sense of smell (anosmia) that may occur by day 3 of a CoV2 infection (Speth et al., 2020), as cells in the nasal cavities support olfactory mucous membranes needed for the perception of smell. Proximity to the brain also suggest that CoV2 infections could impact the brain in ways that other SARS viruses cannot. Moreover, cardiovascular and immunological complications triggered by CoV2 could also be explained on the basis of long-term insult of endothelial cells by viral sequestration of the ACE2 receptor (Gurley and Coffman, 2008).

### pH-switch across species

Further supporting the observation that CoV2 is unique among other coronaviruses is shown in Table 2 that compares sequence alignments of pH domains in RBDs of both CoV1, CoV2, MERS, as well as other closely related zoonotic viruses. CoV2, and related coronaviruses in one species of pangolin and some bats do not share the pH-switch present in CoV1, instead they share the Lys458 stabilization motif. However, these zoonotic viruses still have pH-switches that co-localize next to the pH-switch in the RBD of MERS structure (Fig. 1). Interestingly, different bat-infecting strains show putative pH-switches that are closer in both sequence and structure to either MERS or pangolin-associated coronavirus. While we have not yet found the species or strain where the loss of the pH-switch first occurred, these relationships point at the possible zoonotic origin of CoV2 as well as evolutionary pressures to preserve the pH-switch. It is noteworthy that DPP4, the receptor of the MERS RBD, is not found in nasal epithelial cells (Meyerholz et al., 2016).

**Table 2.**
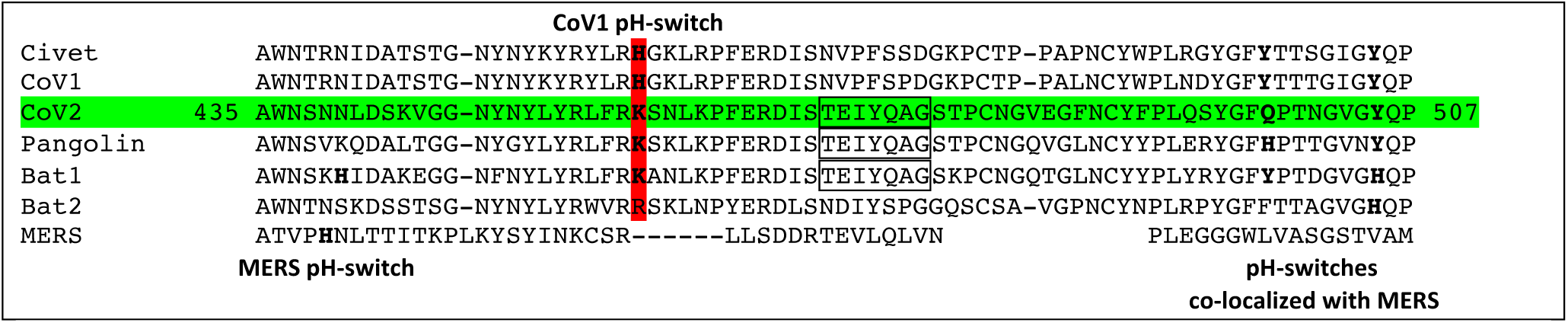
Sequence alignments of pH domains in RBD CoV2 and related viruses. The His*→*Lys mutation is highlighted in red. Conserved motif relevant for interaction with K458 is enclosed in a rectangle for Cov2, Pangolin and Bat1. NCBI Locus AAU04646 (Civet); 6CRV_C (CoV1); 6VSB_A (CoV2); QIQ54048 (Pangolin); QHR63300 (Bat1); AGZ48806 (Bat2); QBM11748 (MERS).

### Outlook

This newly discovered difference in protein sequence in the receptor binding domain of the spike glycoprotein and its impact on receptor binding reveals a mechanism that allows SARS-CoV2 internalization to take advantage of the high expression of ACE2 in the nasal epithelium—resulting in increased retention times in the upper respiratory tract and augmented infectivity. This mechanism reconciles observed epidemiological traits and pathologies specific to SARS-CoV2 and explains differences with those associated with SARS-CoV, which due to its stick-and-slip pH-switch is unable to efficiently undergo endocytosis in the nasal cavity.

SARS-CoV also has a higher infected-fatality rate than SARS-CoV2. While the evolutionary advantage of higher infectivity by SARS-CoV2 in the nasal area is clear, this property comes at the expense of an important regulatory mechanism that would have allowed this virus to more readily move in other organs and tissues. In fact, the life-cycle of SARS-CoV2 is significantly slower than that of SARS-CoV because CoV2 is essentially immobilized at its initial cell receptor contact. Thus, it seems unlikely that the diffusion limited recruitment of ACE2 receptors affecting the virus in the respiratory airways would also be the limiting step in tissues.

After internalization, the virus is encapsulated in a vesicle supported by RBD/ACE2 complexes. The actual final number of complexes in each vesicle should vary above a given threshold, though not much is known about the details of this process. Contrary to SARS-CoV, CoV2 complexes would be expected to have greater difficulty slipping and breaking. It is not difficult to imagine that for vesicles compressed by an excess of receptors the fusion with the early endosome might be hindered, hosting a population of viruses that could stay latent or activate at much later times. This simple mechanism could underlie the still anecdotal evidence for infection recurrence (Chen et al., 2020), as well as extremely long-term of viral shedding.

Collectively, our studies provide insight pertinent to the molecular basis of viral infectivity and, at the same time, validate this form of thermodynamic and molecular modeling as an approach to probe the evolution of the next SARS-mediated pandemic. From a therapeutic perspective, our findings linking viral pathology with long-term viral infection/retention in nasal epithelium of the upper respiratory tract suggest that vaccine development should not just concentrate on fighting systemic infection through induction of IgG responses, but should instead aim to elicit high titers of secretory IgA antibodies capable of neutralizing the virus in the nasal mucosa. Therefore, intranasal delivery of a vaccine with strong IgA producing potential is a logical approach to consider as the next step in countering the current and future pandemics that, like SARS-CoV2, export evolutionarily successful strains via higher transmission rates.

## Acknowledgements

This work was supported by NIH GM097082, NS043277 and T32EB009403. CJC is grateful to Dr. Jose Padial for his inspiration in starting this project, to Dr. Dana Ascherman for helping in the immunobiology component of the model, to Drs. Marc Herant and Harinder Singh for successfully challenging earlier interpretations of the model, to Dr. AR Carvunis for highlighting evolution, and to Dr. Olja Finn for her enthusiasm for our model and helpful suggestions.

## Author Contributions

CJC developed the mechanistic ideas and structural modeling, and wrote the paper. KAP and US performed and analyzed the molecular dynamics simulations.

## Declaration of Interests

The authors declare no competing interests.

## Supplementary Information

## Methods

Atomic coordinates for starting structures were acquired from the Protein Data Bank [1]: 2DD8 was used for CoV1 RBD and 6LZG was used for the CoV2 RBD. The RBD from 2DD8 (bound to neutralizing antibody) was chosen instead of that in PDB ID 2AJF (bound to ACE2) as a starting structure as it includes otherwise missing portions of the domain. Modification of His to different tautomeric or protonation states was done with PyMol’s Mutagenesis Wizard [2]. Molecular dynamics simulations (MDs) were carried out with pmemd.cuda from AMBER18 [3-5] using AMBER ff14SB force field [6] and Generalized Amber Force Field (GAFF) [7]. We used tLeap binary (part of AMBER18) for solvating the structures in a cubed TIP3P water box with a 10 Å distance from structure surface to the box edges, and closeness parameter of 0.75 Å. The system was neutralized and solvated. Simulations were carried out after minimizing the system, gradually heating the system from 0 K to 3 00K over 50 ps, and equilibrating the system for 1 ns at NPT. 500 ns of production was then carried out using NPT at 300 K with the Langevin thermostat, a non-bonded interaction cut off of 8 Å, time step of 2 fs, and the SHAKE algorithm to constrain all bonds involving hydrogens.

Clustering was completed using CPPTRAJ [8] and H-bond and RMSD calculations were done with VMD [9].

All figures were drawn using PyMOL [2] and GNUPLOT.

---

**Figure S1.**
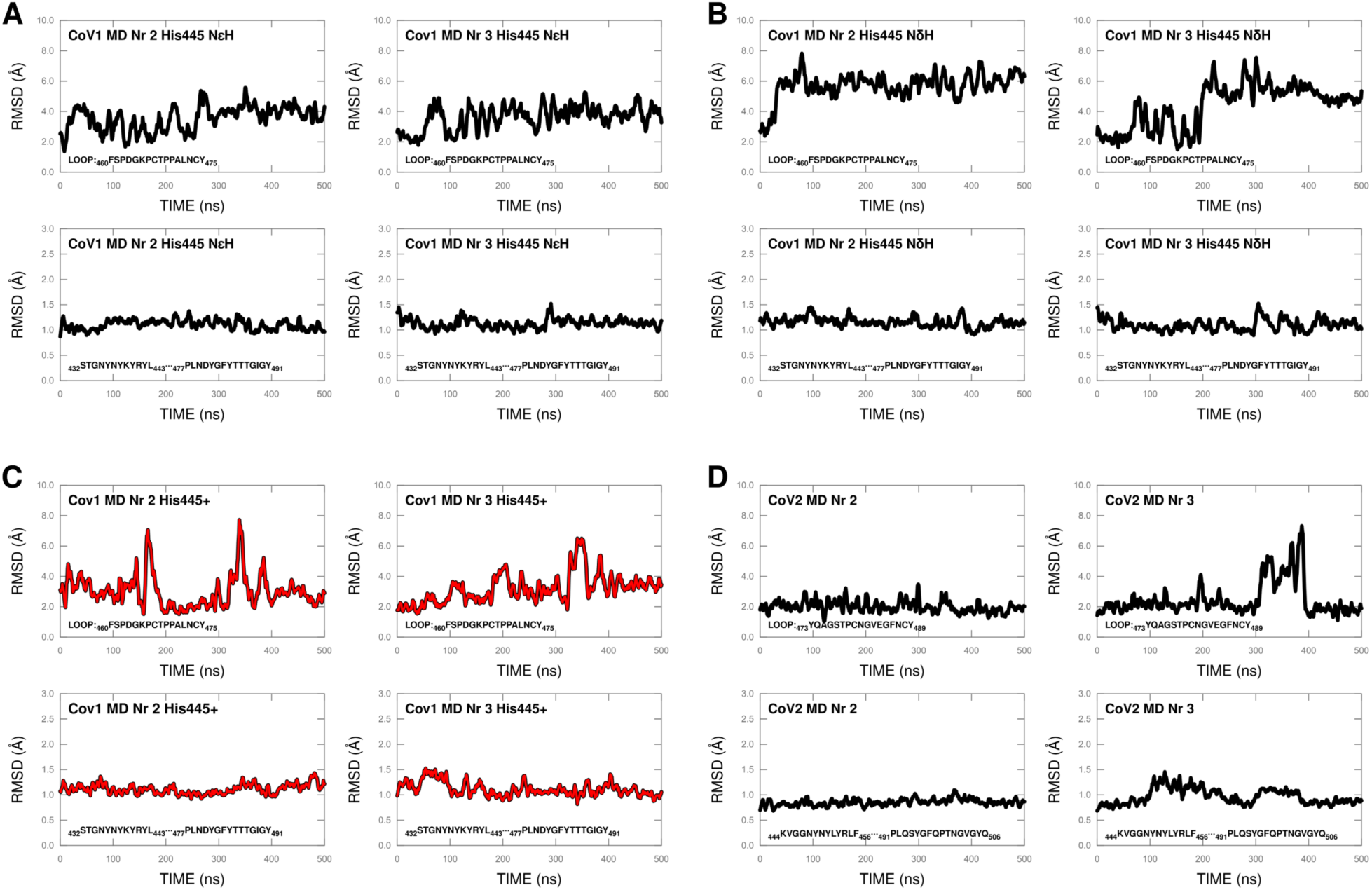
Dynamics of binding interface – additional MDs. Switching Loop and remainder of the binding interface (not including Loop residues) RMSD relative to co-crystal structures as a function of time as described in Fig 3. **A**. Two independent MDs of CoV1 with unprotonated H445 and hydrogen on Nε. Top: RMSD of Loop shows unstable not-bound-like conformations. Bottom: RMSD of the remainder of the binding interface show consistent bound-like phenotype. **B**. Two extra MDs of CoV1 with unprotonated H445 and hydrogen on Nδ. Top: RMSD of Loop shows not-bound-like phenotype. Bottom: RMSD of the remainder of the binding interface show stable bound-like conformations. **C**. Two additional MDs of CoV1 with protonated H445^+^ shows significant bound-like behavior (RMSD ∼ 2-3 Å) for Loop (Top); and, RMSD ∼ 1 Å for binding interface without the loop (Bottom). **D**. Two additional MDs of CoV2: RMSD as a function of time shows consistent bound-like behavior for both the Loop region homologous to CoV1 (Top), and the remainder of the binding interface (Bottom).

**Figure S2.**
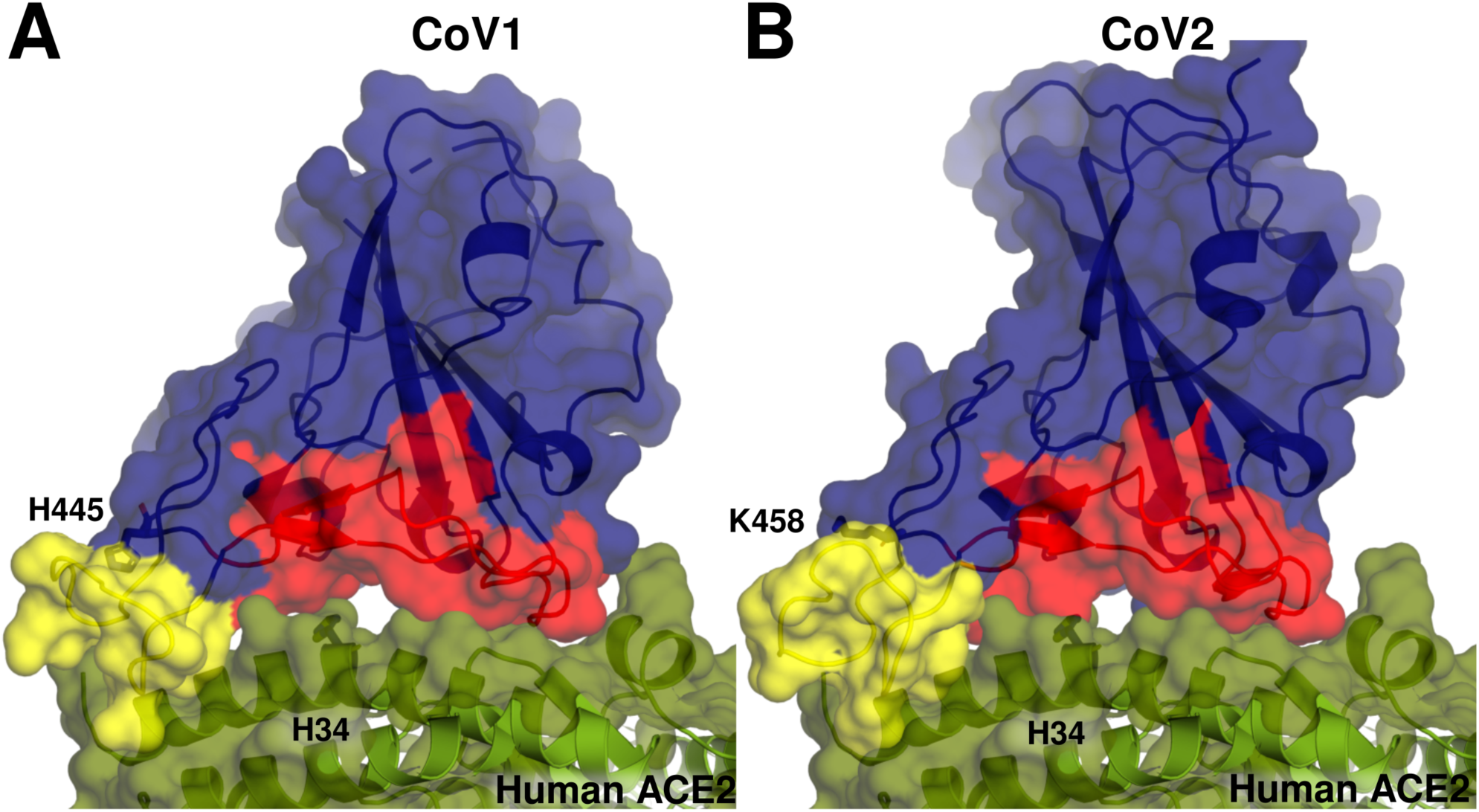
“Door-knob” binding interface of CoV1 (PDB 2AJF) and CoV2 (PDB 6LZG). Surface representation of the co-crystals reveal two characteristic lobes with flat and mostly superficial contacts. Yellow surface corresponds to pH-switch loop and red surface indicates remaining of binding interface.

## Notes

### Competing Interest Statement

The authors have declared no competing interest.

